# Peripheral voltage-gated calcium channels in skin are essential for transient neurogenic thermal hyperalgesia in mice

**DOI:** 10.1101/2020.07.28.225615

**Authors:** Daniel M DuBreuil, Eduardo Javier Lopez Soto, Daniel Li, Diane Lipscombe

## Abstract

Voltage-gated Ca_V_2.2 calcium channels are expressed in nociceptors, at pre-synaptic terminals, soma, and axons. Ca_V_2.2 channel inhibitors applied to the spinal cord relieve pain in humans and rodents, especially during pathological pain, but a biological function of nociceptor Ca_V_2.2 channels in processing of nociception, outside pre-synaptic terminals, is not explored. Here, we demonstrate that functional Ca_V_2.2 channels in skin are required for thermal hyperalgesia following intraplantar capsaicin exposure. We provide evidence that Ca_V_2.2 channels at nociceptor free endings release inflammatory signals, ATP and IL-1β. We assess the role of Ca_V_2.2 splice isoforms to capsaicin-induced hyperalgesia measured by thermal and mechanical stimuli. Our data reveal a critical role for peripheral Ca_V_2.2 channels in skin in neurogenic thermal hyperalgesia but not in mechanical hypersensitivity. Inhibition, or the complete lack, of peripheral Ca_V_2.2 channels blunts the hyperalgesia response *in vivo*.

## INTRODUCTION

Voltage-gated Ca_V_2.2 (N-type) and Ca_V_2.1 (P-type) calcium channels are considered the primary sources of calcium that elicits neurotransmitter release from nociceptor terminals in the spinal cord dorsal horn.^1^ Opiate analgesics inhibit presynaptic Ca_v_2.2 channels in the spinal cord dorsal horn to reduce Ca_V_2.2 channel activity and decrease synaptic transmission.^2,3^ Ca_V_2.2 channels in nociceptors are also targets of non-opioid analgesics such as the Ca_V_2.2 channel inhibitor ω-conotoxin MVIIA (Prialt®, Ziconotide, or SNX-111).^4,5^ ω-Conotoxin MVIIA is effective against chronic, intractable pain in humans and in animal models of inflammatory and nerve injury-induced pain, and it has less effect on acute pain responses.^2,3,5–9^ However, their action on Ca_V_2.2 channels outside of the nociception pathway limit their therapeutic application. Analgesics that preferentially relieve pathological pain, relative to normal nociceptive pain, are highly attractive from a therapeutic perspective, but current dogma that Ca_V_2.2 channels at dorsal horn central synapses are the critical targets – does not account fully for their preferential action on pathological pain states.

Ca_V_2.2 channels are expressed throughout nociceptors at presynaptic terminals, soma, and peripheral structures, including the sciatic nerve, but their biological significance at sites beyond a well-accepted role at presynaptic terminals, is not well studied or understood.^1,10–15^ Nociceptor free nerve endings in skin release inflammatory signaling molecules that contribute to inflammatory hyperalgesia.^12,16–22^ For example, neuropeptides calcitonin gene-related peptide (CGRP) and Substance P released from nociceptor free nerve endings interact with the vasculature to regulate vasodilation and plasma extravasation during inflammation^16^ and released ATP binds to purinergic receptors on neurons, vascular endothelial cells, and immune cells to promote hyperalgesia.^17–19^ The mechanisms that mediate the release of these pro-inflammatory signals are not determined and a role for Ca_V_2.2 channels has not been thoroughly explored ^23^, but the anti-inflammatory action of dual Na_V_1.7 and Ca_V_2.2 channel inhibitors indicate their contributions to peripheral release of pro-inflammatory mediators.^24^

Ca_V_2.2 channel α1 subunits are encoded by the *Cacna1b* gene which generates a subset of possible splice isoforms depending on tissue and cell-type, developmental age, and disease state, and with different functional characteristics.^25–27^ *Trpv1*-expressing nociceptors express Ca_V_2.2 splice isoforms that contain at least two forms, one contains an ubiquitous exon e37b (Ca_V_2.2-e37b) and the other e37a (Ca_V_2.2-e37a) with limited expression across the nervous system.^27^ Ca_V_2.2-e37a channels are trafficked to the cell surface with greater efficiency, and are more sensitive to modulation by morphine compared with Ca_V_2.2-e37b.^14,27–29^ Ca_V_2.2-e37a channel expression is tightly controlled in *Trpv1*-nociceptors by cell-specific, exon-specific hypomethylation. Exon hypomethylation allows binding of the ubiquitous 11-zinc finger CCCTC binding factor (CTCF), which in turn promotes exon recognition and its inclusion in mRNA.^30^ The use of splice isoform specific siRNAs suggest that Ca_V_2.2-e37a channels are particularly critical for some forms of hyperalgesia, although Ca_V_2.2 splice isoform contribute equally to acute nociception.^31^

Here, we explore a role for Ca_V_2.2 channels in nociceptor free nerve endings in skin and compare the contribution of Ca_V_2.2 channel splice isoforms at peripheral and central sites. We show that Ca_V_2.2 channels at central nociceptor termini contribute to the accurate transmission of information about acute noxious stimuli. In addition, we show for the first time that Ca_V_2.2 channels are required for transient neurogenic thermal hyperalgesia induced by capsaicin, and Ca_V_2.2-e37a isoforms appear to be required for maximum hyperalgesia responses.

## RESULTS

### Ca_V_2.2 channels contribute to acute nocifensive responses

The largest component of the somatic high voltage-activated Ca_V_ channel current in nociceptors originates from the gating of Ca_V_2.2 channels.^11,14,15^ They dominate at presynaptic termini in the spinal cord dorsal horn to trigger transmitter release and support transmission of noxious stimuli.^1^ However, their contribution to acute nocifensive behaviors is incompletely defined.^28,32–34^ Studies of Ca_V_2.2-null mouse strains have yielded inconsistent conclusions in this regard for reasons that remain unclear.^32–34^ Therefore, we used a new *Cacna1b* knockout mouse strain (KO) to quantify the role of Ca_V_2.2 channels in acute nociception applying both conventional and optogenetic stimuli in wild-type (WT) and KO mice (Fig. 1). We used immunoblotting and RNA *in situ* hybridization in dorsal root ganglia (DRG) to confirm the absence of Ca_V_2.2 protein (Fig. 1a) and *Cacna1b* RNA (Fig. 1b) in our KO mouse strain. By RNA *in situ* hybridization using e37a-specific probes, we also show the presence of *Cacna1b*-e37a isoform encoding RNAs in a subset of DRG neurons (Fig. 1b).

**Figure 1.**
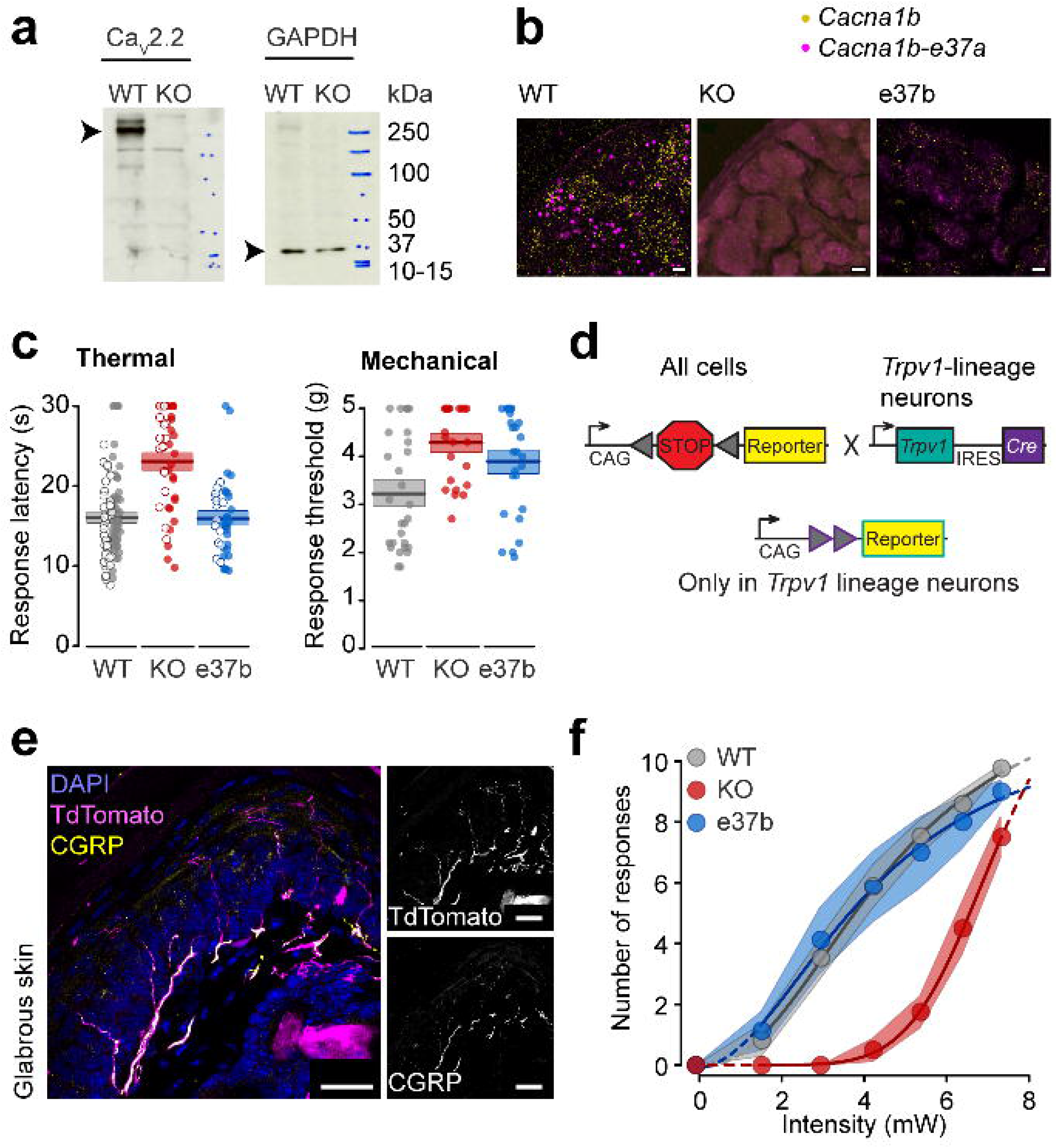
Ca_V_2.2 knockout, but not Ca_V_2.2-e37b only mice exhibit impaired nociception compared to wild type controls. **a.** Western blot shows major full length Ca_V_2.2 α1 subunit protein (~250 kDa band) in whole brain lysate from wildtype mice (WT) and absence in Ca_V_2.2-null mice (KO). Additional minor bands present in WT but not in KO may reflect different isoforms (>250 kDa) or proteolytic fragments (100-150 kDa) of Ca_V_2.2 α1. Additional bands present in both WT and KO brain lysates (~ 150 kDa) reflect non-specific binding of anti-Ca_V_2.2 antibody. GAPDH signals (~ 37 kDa) are present in WT and KO brain lysate in the same blot, stripped and re-probed with anti-GAPDH. **b.** *Cacna1b* mRNA (green) is present in dorsal root ganglia (DRG) of Ca_V_2.2 WT mice, absent in DRG of Ca_V_2.2 KO mice, and present in DRG of Ca_V_2.2-e37b only mice. *Cacna1b-37a* mRNA (red) is only present in dorsal root ganglia (DRG) of Ca_V_2.2 WT mice. Images show amplification of *Cacna1b* pan mRNA probe (green) and *Cacna1b-37a* specific mRNA probe (red). All experiments use RNAScope@ methods, scale bars for all images = 10 μm. **c**. Thermal and mechanical sensitivity compared in Ca_V_2.2 WT (grey), KO (red) and Ca_V_2.2-e37b only (blue) mice, measured by two different experimenters under unblinded (open symbol) and blinded (closed symbol) conditions. Thermal and mechanical sensitivities of KO mice are reduced compared to WT and e37b-only mice. Each value shown represents the median of 3 measurements each from the following number of mice (Thermal: WT n = 146, KO n = 56, e37b n = 42; Mechanical: WT n = 32, KO n = 24, e37b n = 26), together with means (horizontal line) and SEs (shaded area). Mean ± SE were for thermal latency WT: 16.0 ± 0.4 s; KO: 21.8 ± 0.8 s; e37b: 15.9 ± 0.7 s; mechanical threshold WT: 3.35 ± 0.21 g; KO: 4.29 ± 0.16 g; e37b: 3.90 ± 0.21 g. *P* values calculated by univariate ANOVA with Tukey HSD correction for multiple comparisons. Thermal WT vs KO, *p* = 0.0000003; WT vs e37b, *p* = 0.999. Mechanical WT vs KO, p= 0.021; WT vs e37b, *p* = 0.228. with Tukey HSD correction for multiple comparisons. **d**. Scheme showing Cre/loxP breeding strategy to generate heterozygote reporter strains expressing either ChR2 or tdTOMATO (reporter) in *Trpv1*-lineage neurons. We generated nine different strains of mice: WT, KO, and e37b-only in three different JAX mice *Trpv1^tm1(cre)Bbm^/J (Trpv1-Cre;* RRID: IMSR_JAX:017769)^74^, *Gt(ROSA)26Sor^tm32(CAG-COP4^*^H134R/EYFP)Hze^/J (lox-STOP-lox-ChR2-EYFP;* JAX #012569)^75^, and *Gt(ROSA)26Sor^tm14(CAG-tdTomato))Hze^/J (lox-STOP-lox-tdTomato;* RRID: IMSR_JAX:007908)^72^. These nine strains were maintained as homozygotes and, as needed, parents bred for first-generation heterozygote progeny. *Trpv1* is expressed in testes during gamete production so only first-generation progeny were used (see methods). **e**. Free nerve endings of *Trpv1* lineage nociceptors in plantar glabrous skin co-express CGRP. *Left*, overlay shows *Trpv1/tdTomato* (magenta) free nerve endings, anti-CGRP (yellow), and cell nuclei labelled with DAPI (blue). Image is reconstructed from xxx optical sections of xxx um. See methods for details. *Right*, grayscale images of *Trpv1/tdTomato (top)* and anti-CGRP (*bottom*) signals. All scale bars = 25 μm. **f**. Behavioral responses induced by 5 s LED directed to plantar hindpaws of wildtype (WT), KO and e37b-only mouse strains expressing ChR2 in *Trpv1*-lineage neurons (see ***d*** for details). KO mice were less response to 465 nm LED stimulation compare to WT and e37b mouse strains. Each symbol represents the mean ± SE of the number of responses to 10 separate LED stimuli, applied to the following number of mice: WT/Trpv1/ChR2-EYFP, n = 17; *KO/Trpv1/ChR2-EYFP*, n = 9; and *e37b/Trpv1/ChR2-EYFP*, n = 7. Intensity-response curves for each mouse were fit using a 4-parameter logistic function. *P* values for ANOVA with Dunnett T3 correction for multiple comparisons were for WT vs KO: *p* = 0.000002; and WT vs e37b *p* = 0.997.

Paw withdrawal latencies to thermal stimuli were consistently longer in KO mice compared with WT controls (21.8 ± 0.8 s, n = 56 KO mice compared to 16.0 ± 0.4 s, n = 146 WT mice; *p* = 0.0000003). This difference was observed in two independent cohorts of mice assessed by different experimenters and under both blinded and unblinded conditions, with no significant effect of cohort (*p* = 0.089) or interaction between genotype and cohort (*p* = 0.666). KO mice also had higher mechanical paw withdrawal thresholds compared to WT controls (4.29 ± 0.16 g, n = 24 KO mice compared with 3.35 ± 0.21 g, n = 32 WT mice; *p* = 0.021, Fig. 1c). Our results support findings of some^32,34^, but not all^33^ previously reported studies using independently generated KO mice. Ca_V_2.2 channels are important for normal nocifensive responses but, at least in paw withdrawal assays using thermal and mechanical stimuli, other Ca_V_ channels support behavioral responses induced by higher intensity stimuli (Fig. 1c). Our findings are consistent with results from experiments using similar behavioral assays in response to acute pharmacological inhibition of Ca_V_2.2 channels by intrathecal ω-conotoxin MVIIA in wild-type mice^28^ suggesting that there is little or no compensatory increase in the expression of other Ca_V_ channels in the KO mouse strain generated in our lab. This is consistent with studies of other KO mouse strains which have not reported upregulation of non-Ca_V_2.2 Ca_V_ channels.^32–34^

Naturalistic stimuli recruit multiple sensory neuron subtypes. Therefore, to assess the contribution of Ca_V_2.2 channels to behavioral responses following activation of a genetically-defined nociceptor subpopulation, we applied an optogenetic approach using WT and KO mouse strains that express ChR2-EYFP in *Trpv1-lineage* neurons (Fig. 1d-f). In the glabrous skin of the plantar hind paw, where we applied LED light to evoke behavioral responses, *Trpv1-lineage* axon fibers were primarily peptidergic nociceptors based on overlap of *Trpv1-lineage* reporter (TdTomato) with anti-CGRP (Fig. 1e). In both mouse strains, WT and KO, brief exposure of the plantar hind paw to blue LED light elicited robust and stereotyped nocifensive paw withdrawal responses that were light-intensity dependent (Movie 1, Fig. 1f). Compared with WT, KO mice were significantly less responsive to LED stimulation over a wide range of light intensities (WT, n = 17 mice; KO, n = 9 mice; *p* = 0.000002, Fig. 1f). Our analyses demonstrate a clear separation in the behavioral sensitivities of KO and WT mice evoked by activating *Trpv1-lineage* nerve endings in skin. At high LED intensities that activate more nociceptor free nerve endings, the responsiveness of KO and WT mice is similar (Fig. 1f).

We showed previously that Ca_V_2.2 currents in nociceptors reflect the combined activity of multiple splice isoforms generated from the *Cacna1b* gene.^27^ Of particular interest is a pair of mutually exclusive alternatively spliced exons, e37a and e37b. Ca_V_2.2-e37a channels are enriched in *Trpv1*-expressing nociceptors as compared to most other neurons which primarily express Ca_V_2.2-e37b channels.^27,30^ We used knock-in mice that only express Ca_V_2.2-e37b channels (e37b, Fig. 1b)^14,28^ to show that e37b is able to substitute for e37a and support normal thermal and mechanical sensitivity (e37b, thermal: 15.9 ± 0.7 s, n = 42 mice, *p* = 0.999 compared to WT; e37b, mechanical: 3.90 ± 0.21 s, n = 26 mice, *p* = 0.228 compared to WT, Fig. 1c). We also expressed ChR2-EYFP in *Trpv1*-lineage neurons of e37b mice and found LED-induced nocifensive behavior to direct nociceptor activation was not distinguishable from WT (e37b, n = 7 mice, *p* = 0.997, Fig. 1f), consistent with our data using traditional nociception assays.^14^ For example, in previous studies using both Ca_V_2.2-e37a only and Ca_V_2.2-e37b only mouse strains, we demonstrated that the basal thermal responsiveness of mice is not impacted by selective removal of either exon.^14,30^ Below, we compare the relative importance of Ca_V_2.2-e37a and Ca_V_2.2-e37b channels to mediate synaptic transmission in the spinal cord dorsal horn.

### Postsynaptic currents in spinal cord dorsal horn neurons elicited by optical stimulation are smaller in Ca_V_2.2-null mice compared to wild-type

We recorded light-activated post-synaptic currents in lamina II neurons of spinal cord dorsal horn in acute slices from *WT/Trpv1*^Chr2EYFP+/-^ (WT), *KO*/*Trpv1*^Chr2EYFP+/-^ (KO), and *e37b*/*Trpv1*^Chr2EYFP+/-^ (e37b) mice to test the hypothesis that the complete absence of Ca_V_2.2 channels reduces synaptic transmission from nociceptor pre-synaptic terminals (Fig. 2a, 2b). *Trpv1*-lineage afferent fibers terminate in laminae I, II, and III, as demonstrated by the expression of ChR2-EYFP in areas both superficial to and deeper than IB4 binding (Fig. 2a). The total charge and amplitude of light-evoked postsynaptic currents (PSCs) were consistently smaller in neurons from KO mice compared to those in WT controls (KO amplitude: −252.1 ± 48.2 pA, n = 12 cells; charge: −7.2 ± 1.1 pC, n = 12 cells; WT amplitude: −511.8 ± 55.1, n = 20 cells, *p* = 0.008; charge: −18.6 ± 1.7 pC, n = 20 cells, *p* = 0.000051; Fig. 2c and 2d). Light-evoked PSCs in KO neurons decayed more rapidly compared with WT as reflected by a more transient, and smaller, slow component of the light-evoked PSC (WT τ_weighted_ = 42.3 ± 4.0 ms, n = 20 cells; KO τ_weighted_ = 27.4 ± 2.3 ms, n = 12 cells; *p* = 0.009, Fig 2e; WT τ_slow_ = 119.0 ± 8.9 ms, 28.3 ± 2.6% amplitude, n = 20 cells; KO τ_slow_ = 93.1 ± 8.4 ms, 22.0 ± 2.8% amplitude, n = 12 cells; *p* (τ_slow_) = 0.122, *p* (amplitude) = 0.282, Fig. 2f, 2h). The uniformly smaller synaptic responses in spinal cord slices from KO mice are consistent with the attenuated LED-evoked behavioral responses (Fig. 1f) and also suggest little or no compensation from other Ca_V_ channels.

**Figure 2.**
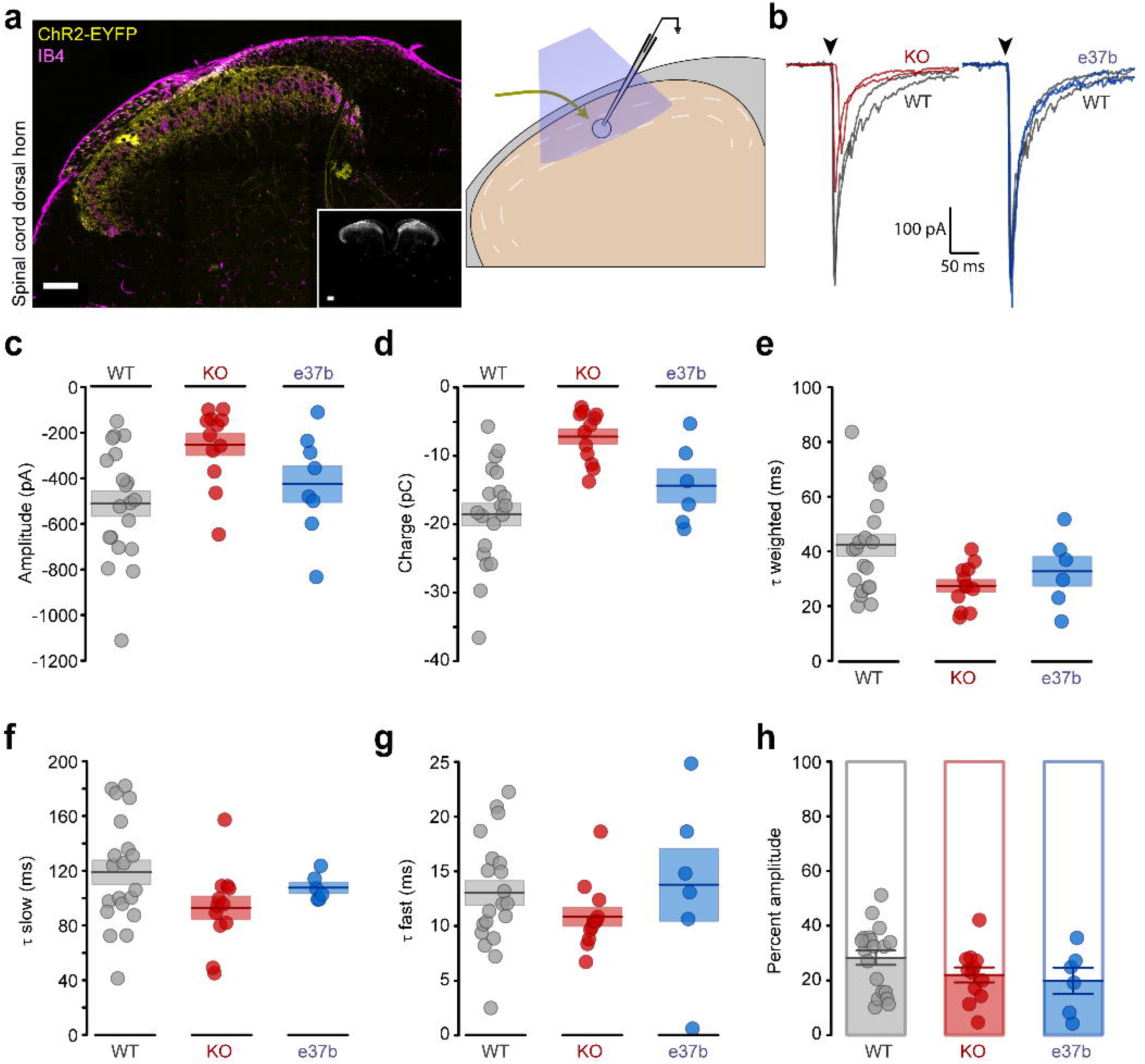
Synaptic currents recorded in spinal cord slices from WT, KO and e37b mice. **a**. *Left:* Transverse section of lumbar spinal cord dorsal horn from WT/Trpv1/ChR2-EYFP mouse showing ChR2-EYFP (yellow) and IB4 (magenta) to identify lamina II. Inset: grayscale image of whole spinal cord section from WT/Trpv1/ChR2-EYFP mouse showing reporter expression is restricted to dorsal horns. Scale bars represent 100 μm. *Right:* Schematic of experimental setup for recording light activated synaptic currents in postsynaptic lamina II neurons. **b**. Synaptic currents in slices from WT/Trpv1/ChR2-EYFP mice (grey), KO/Trpv1/ChR2-EYFP (red, *left*), and e37b/Trpv1/ChR2-EYFP (blue, *right*
) mice. **c-h**) Comparison of peak current amplitudes (**c**), total charge (d), weighted time constant (**e**), slow time constant (**f**), fast time constant (**g**), and component contributions to response amplitude (**h**) for synaptic currents. For **c-g**, symbols represent values of individual neurons, together with mean (horizontal line) ± SE (shaded area). For **h**, symbols represent the relative contributions of fast (shaded) and slow (white) components of the synaptic currents measured from each cell together with mean ± SE. Analyses summarize properties of synaptic currents recorded from 20 WT, 12 KO, and 6 e37b neurons. Mean ± SE and p values assessed by multivariate ANOVA were for total charge: WT: −20.6 ± 2.5 pC, KO: −7.2 ± 1.1 pC, *p* = 0.000051 (KO vs WT); e37b: −14.3 ± 2.5 pC, *p* = 0.328 (e37b vs WT); peak amplitude: WT: −547.6 ± 76.0 pA, KO: −252.1 ± 48.2 pA, *p* = 0.008 (KO vs WT), e37b: −425.3 ± 80.7 pA, *p* = 0.950 (e37b vs WT); weighted time constant: WT: 44.5 ± 5.9 ms, KO: 27.4 ± 2.3 ms, *p* = 0.009 (KO vs WT), e37b: 32.7 ± 5.4 ms, *p* = 0.431(e37b vs WT); slow time constant: WT: 123.6 ± 12.9 ms, KO: 93.1 ± 8.4 ms, *p* = 0.122 (KO vs WT), e37b: 107.6 ± 4.0 ms, *p* = 0.577 vs WT; fast time constant: WT: 11.7 ± 1.6 ms, Ca_V_2.2-null: 10.8 ± 0.9 ms, *p* = 0.351 (KO vs WT), e37b: 13.8 ± 3.3 ms, *p* = 0.995 (e37b vs WT); contribution of slow component to response amplitude: WT: 29.9 ± 3.0%, KO 22.0 ± 2.8%, *p* = 0.282 (KO vs WT), e37b: 19.9 ± 4.8%, *p* = 0.922 (e37b vs WT).

By contrast, light-evoked PSCs in acute slices from e37b mice were not consistently different from WT with respect to amplitude, total charge, and decay time (e37b amplitude: −425.3 ± 80.7 pA, *p* = 0.950; charge: −14.3 ± 2.5 pC, *p* = 0.328; decay time: *τ* = 32.7 ± 5.4 ms, *p* = 0.431; n = 8 cells; Fig. 2c-h). This parallels the normal nocifensive responses observed in e37b only mice compared with WT (Fig. 1f).

We measured post-synaptic currents during repetitive stimulation of *Trpv1-lineage* nociceptors to assess the role of Ca_V_2.2 channels in responses to bursts of action potentials that occur during natural stimulation. Differences in the degree of depression of post-synaptic current amplitudes, during repetitive stimulation, can also be indicative of pre-synaptic involvement.^35–37^ We applied 10 light pulses at 1 Hz to acute spinal cord slices from WT, KO, and e37b mice (Fig. 3a). In all three genotypes, post-synaptic currents followed 1 Hz stimulation perfectly, with no synaptic failures (Fig 3b), but the degree of depression at KO synapses was consistently reduced as compared to WT and e37b synaptic currents (KO: 86% of the initial EPSC amplitude compared to 51% and 54% in WT and e37b, respectively; WT vs e37b: *p* = 0.940; WT vs KO: *p* = 0.001; Fig 3c). These results are consistent with reduced neurotransmitter release probability at nociceptor synapses lacking Ca_V_2.2 channels as compared to WT.

**Figure 3.**
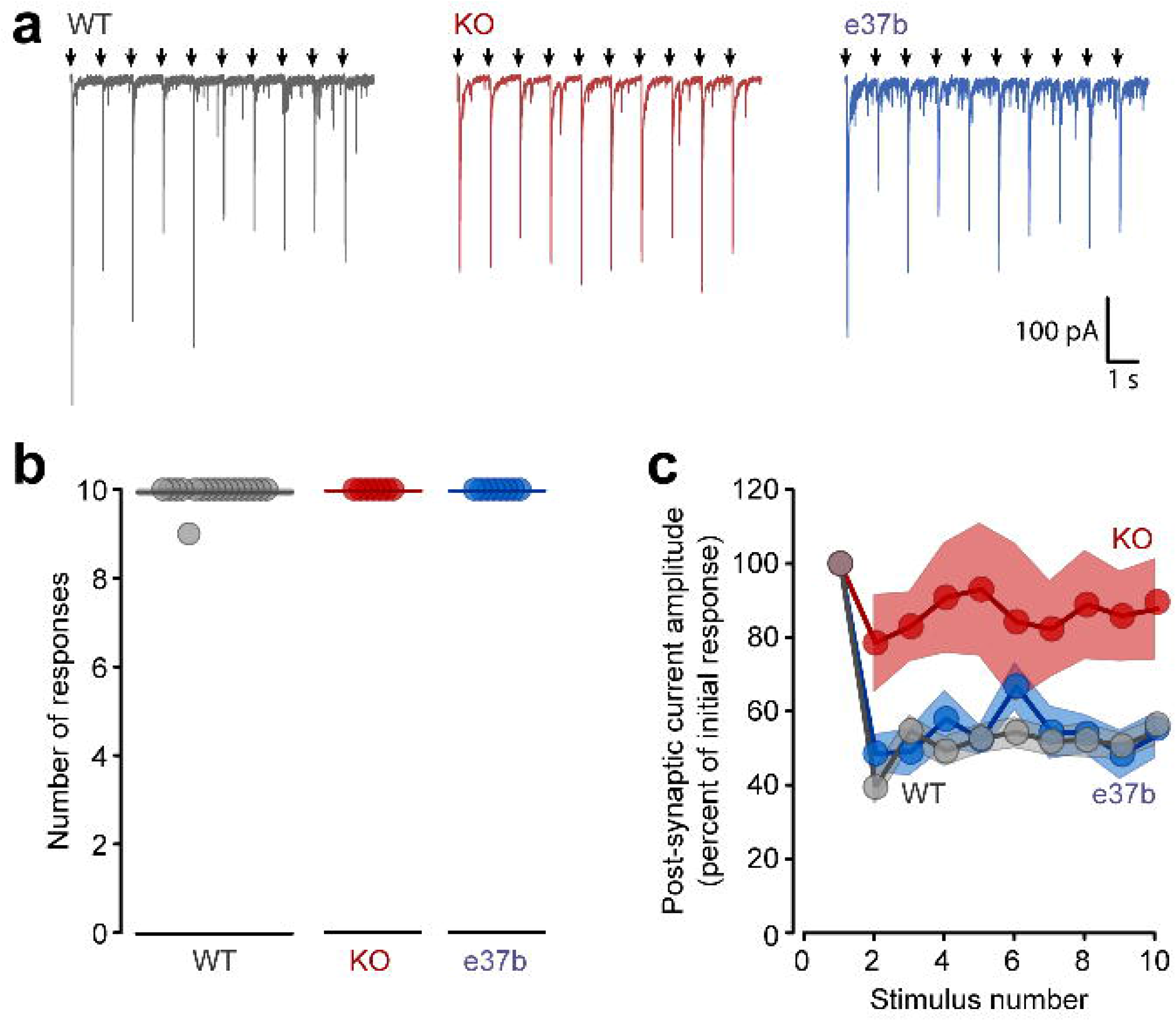
Temporal features of synaptic currents in WT, KO and e37b mice. **a**. Synaptic currents measured from postsynaptic neurons elicited by a series of 10 optical stimuli applied at 1 Hz, in spinal cord slices from WT/Trpv1/ChR2-EYFP (*left*, grey), KO/Trpv1/ChR2-EYFP (*middle*, red), and e37b/Trpv1/ChR2-EYFP (*right*, blue) mice. **b**. Number of synaptic currents elicited by 1 Hz optical stimulation recorded in spinal cord splices from WT (n = 17), KO (n = 7) and e37b (n = 8). Each symbol represents recordings from an individual slice. **c**) Relative amplitude of postsynaptic currents compared to the initial response, evoked by optical stimulation at 1 Hz for WT (grey, n = 9 slices), KO (red, n = 7 slices) and e37b (blue, n = 8 slices) mice. *P* values assessed by repeated measures ANOVA and Tukey HSD correction for multiple comparisons were for WT vs KO, *p* = 0.001; WT vs e37b, *p* = 0.940.

To this point, we show that Ca_V_2.2 channels are important for normal behavioral sensitivity and contribute to transmission of information about noxious stimuli. At higher stimulus intensities, the recruitment of additional nerve fibers likely compensates for overall reduced release probability in KO mice. Ca_V_2.2 channels, and Ca_V_2.2-e37a channel isoforms in particular, are preferentially implicated in the development of several pain states including hyperalgesia.^2,6,31^ We therefore explored the role of Ca_V_2.2 in a robust model of transient hyperalgesia.

### Ca_V_2.2 channels are critical for capsaicin-induced thermal hyperalgesia

Capsaicin acts specifically on neuronal TRPV1 channels to induce hyperalgesia and intraplantar capsaicin is a classic model of neurogenic inflammation.^38,39^ We used intraplantar injections of capsaicin (0.1% w/v) to depolarize nociceptors and, at 15 and 30 minutes after injection, measured thermal and mechanical hyperalgesia in WT, KO and e37b mice (Fig. 4a). Two experimenters, using unblinded and blinded protocols, tested two independent cohorts of mice. In the unblinded condition, only capsaicin was injected, whereas in the blinded condition either capsaicin or saline was injected into WT and KO mice. For both unblinded and blinded experiments, intraplantar capsaicin elicited thermal hyperalgesia in all WT mice tested (unblinded: n = 10 mice; blinded: n = 23 mice) and injection of saline in WT mice had no effect (blinded: n = 6 mice). Because of the different experimental conditions, as well as a significant interaction between genotype and cohort (*p* = 0.036), we analyzed blinded and unblinded experiments independently.

**Figure 4.**
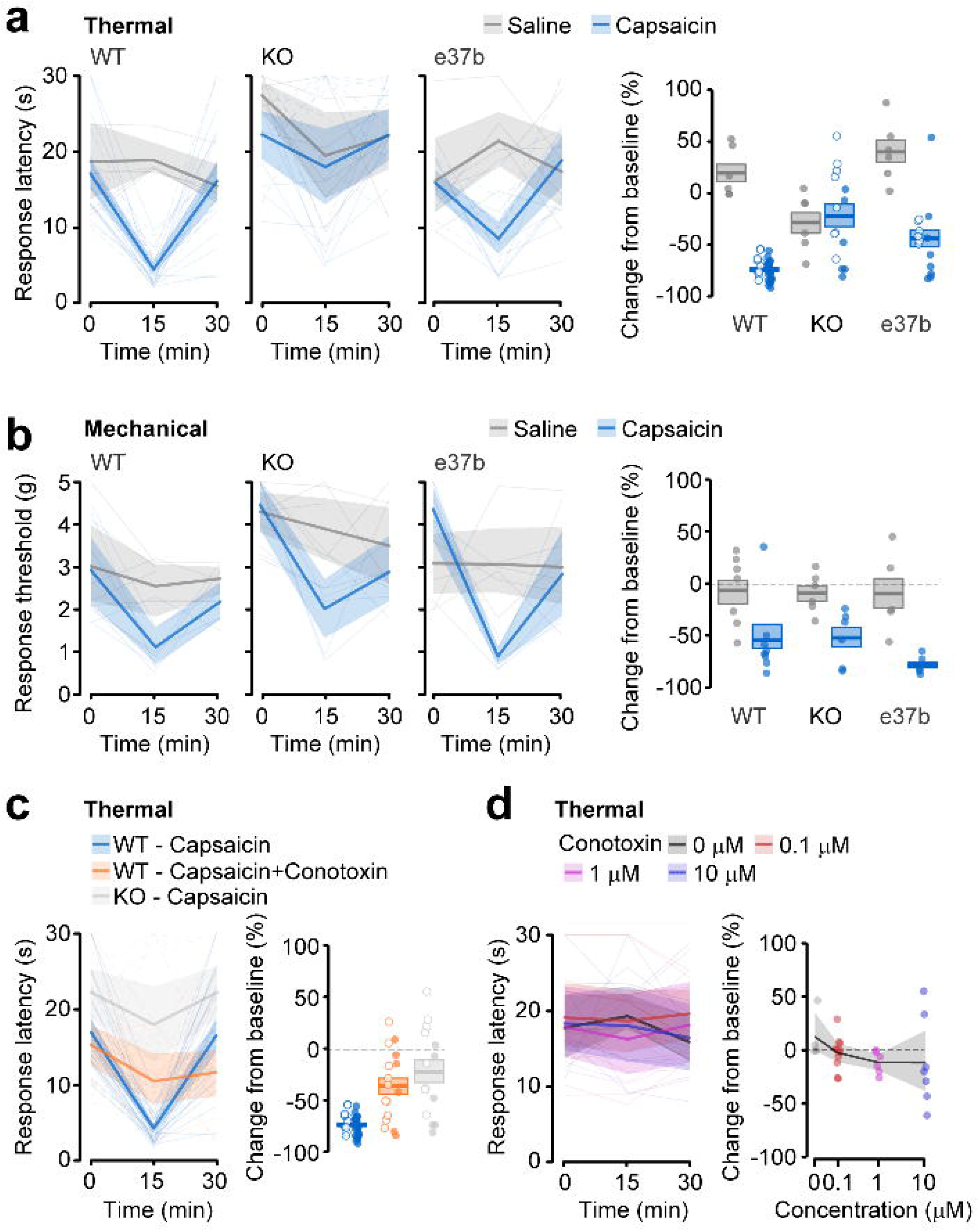
Capsaicin-induced thermal hyperalgesia requires peripheral Ca_V_2.2 channels in skin. **a**. Thermal sensitivity of plantar hind paw of WT, KO, and e37b mice measured before (0 min), and 15 min and 30 mins after injection of 0.1% w/v capsaicin (*blue*) or saline (*grey*). Experiments were conducted in two independent cohorts, by two different experimenters, under blinded (solid lines, open symbols) and non-blinded conditions (dashed lines, open symbols). *Left*, responses from individual mice (thin lines, dashed and solid), average data (thick lines), and SE (shaded areas) are shown for all conditions. *Right*, thermal responses represented as change from baseline for individual mice (symbols, solid and open), average (horizontal line), and SE (shaded areas). Mean and SE were for WT/Saline: n = 6 mice, 19.7 ± 9.8%; WT/Capsaicin: n = 33 mice, −74.0 ± 1.8%; KO/Saline: n = 6 mice, −28.3 ± 11.4%; KO/Capsaicin: n = 15 mice, −12.5 ± 14.6%; e37b/Saline: n = 6 mice, 40.1 ± 12.1%; e37b/Capsaicin: n = 15 mice, −43.5 ± 8.7%. *P* values calculated by univariate ANOVA and Tukey HSD correction for multiple comparisons using pooled data from blinded and unblinded experiments were……. **b**. Mechanical sensitivity of plantar hind paw of WT, KO, and e37b mice measured before (0 min), and 15 min and 30 mins after injection of capsaicin (*blue*) or saline (*grey*). All experiments were performed under blinded conditions. Mean and SE were for WT/Saline: n = 7 mice,-6.9 ± 12.8%; WT/Capsaicin: n = 8 mice, −54.7 ± 13.4%; KO/Saline: n = 6 mice, −9.4 ± 7.8%; KO/Capsaicin: n = 6 mice, −52.2 ± 10.5%; e37b/Saline: n = 6 mice, 16.1 ± 29.6%; e37b/Capsaicin: n = 6 mice, −78.3 ± 3.4%. *P* values calculated by univariate ANOVA and Tukey HSD correction for multiple comparisons were: WT v KO, *p* = 0.994; WT vs e37b, *p* = 0.996. **c**. Thermal sensitivity of plantar hind paw of WT and KO mice measured before (0 min), and 15 min and 30 mins after injection of 0.1% w/v capsaicin in WT (*blue*, n = 33, same data as in **a**), 0.1% w/v capsaicin + 1 μM ω-conotoxin MVIIA in WT (*orange*, n = 17), and 0.1% w/v capsaicin in KO (*gray*, n = 15, same data as in **a**). Experiments were conducted in two independent cohorts, by two different experimenters, under blinded (solid lines, open symbols) and nonblinded conditions (dashed lines, open symbols). *Left*, responses from individual mice (thin lines, dashed and solid), average data (thick lines), and SE (shaded areas) are shown for all conditions. *Right*, thermal responses represented as change from baseline for individual mice (symbols, solid and open), average (horizontal line), and SE (shaded areas). Mean and SE percent change were −18.7 ± 19.3%, *p* = 0.003 (WT con vs WT ω-conotoxin MVIIA). **d**. Thermal sensitivity of plantar hind paw of WT mice measured before (0 min), and 15 min and 30 mins after injection of different concentrations of ω-conotoxin MVIIA (0, 0.1, 1, 10 μM). P value calculated by univariate ANOVA for main effect of dose was *p* = 0.835. For all plots, symbols are values measured from individual animals, average values (solid lines) and SE (shaded areas).

In unblinded conditions, we directly compared the effect of capsaicin in WT and KO mice and observed reduced thermal hypersensitivity following capsaicin injection in KO mice relative to WT (change in response latency following injection: WT: −69.2 ± 3.3%, n = 10 mice; KO: 10.1 ± 19.2%, n = 9 mice; *p* = 0.002). In blinded conditions, we compared the effects of capsaicin and saline within each genotype, to control for genotype-dependent effects of intraplantar injection itself. WT mice exhibited thermal hypersensitivity following capsaicin, but not saline (WT mice-saline: 19.7 ± 9.8% change in response latency from baseline, n = 6; capsaicin: −76.1 ± 2.0% change in response latency from baseline, n = 23; *p* < 0.0000001), whereas responses of KO mice to saline and capsaicin injections were not consistently different. These data show that responses to intraplantar capsaicin and saline were not distinguishable in KO mice (KO mice-saline: −28.3 ± 11.4% change in response latency from baseline, n = 6; capsaicin: −46.4 ± 15.0% change in response latency from baseline, n = 6; *p* = 0.846).

In two independent cohorts of animals assessed by two investigators, capsaicin-induced thermal hyperalgesia was substantially reduced in KO animals as compared to WT. By contrast, capsaicin-induced mechanical hypersensitivity was not distinguishable between WT and KO animals (WT mice-capsaicin: −54.7 ± 13.4% change in response threshold from baseline, n = 8; KO mice-capsaicin: −52.2 ± 10.5% change in response threshold from baseline, n = 6; *p* = 0.999). We conclude that Ca_V_2.2 channels are critical to support capsaicin-induced transient thermal hyperalgesia, but not transient mechanical hyperalgesia, at least under the conditions of our experiments (Fig. 4b).

We next assessed capsaicin-mediated thermal and mechanical hyperalgesia in e37b mice and found reduced thermal hyperalgesia relative to WT (e37b −43.5 ± 8.7% decrease in response latency from baseline; 42.8 ± 11.7% decrease in effect size; *p* = 0.018, n = 15 mice, Fig. 4a), with no significant effect on mechanical hyperalgesia (e37b capsaicin: −78.3 ± 3.36% change in response threshold from baseline, n = 6; *p* = 0.862). In all three mouse strains, robust, spontaneous behavioral responses developed immediately following capsaicin injection and lasted for approximately 5 minutes. This indicated successful injection of active capsaicin. These results point to a new role for Ca_V_2.2 channels in capsaicin-induced thermal hyperalgesia via TRPV1-responsive nociceptors. Our data also suggest that Ca_V_2.2-e37a channels may be necessary for the full thermal hyperalgesia response to capsaicin.

Although we applied capsaicin locally (intraplantar), and hyperalgesia developed rapidly in WT mice, our experiments using KO mice do not rule out contributions from Ca_V_2.2 channels located at sites outside of the skin to capsaicin-induced hyperalgesia.

### Ca_V_2.2 channel activation at peripheral sites is necessary for hyperalgesia

To test if local activation of Ca_V_2.2 channels in skin is essential for capsaicin-induced hyperalgesia, we used local pharmacological inhibition of Ca_V_2.2 channels by ω-conotoxin MVIIA. Capsaicin (0.1% w/v) and ω-conotoxin MVIIA (1 μM) were co-injected into the plantar hind paw of 17 WT mice (Fig. 4c) by two different experimenters, under blinded and unblinded conditions, in two independent mouse cohorts. Injection volumes were consistent across capsaicin/ω-conotoxin MVIIA mixture or capsaicin alone, and robust, spontaneous behavioral responses resulting from direct nociceptor activation was observed in each mouse receiving the injection. Capsaicin-induced thermal hyperalgesia was reduced by ω-conotoxin MVIIA relative to injection of capsaicin alone (−18.7 ± 19.3% change in response latency from baseline; 75.4 ± 25.6% decrease in effect size; *p* = 0.003, Fig. 4c). There was no significant effect of cohort (*p* = 0.697) or interaction between injection type and cohort (*p* = 0.495), thus all animals were pooled for analysis.

The magnitude of the blocking action of intraplantar ω-conotoxin MVIIA on capsaicin-induced thermal hyperalgesia in WT mice (75.4 ± 25.6% decrease in effect size) was similar to the reduction in capsaicin-induced thermal hyperalgesia in KO mice compared to WT (83.1 ± 20.0% decrease in effect size). This provides additional support that Ca_V_2.2 channels in skin are a major contributor to the development of capsaicin-induced thermal hyperalgesia.

Intrathecal ω-conotoxin MVIIA attenuates basal nociception,^8,28^ therefore, to control for the unlikely possibility that intraplantar ω-conotoxin MVIIA reaches the spinal cord thereby confounding our interpretation, we assessed behavioral responses in WT mice to thermal stimuli in the presence of increasing concentrations of ω-conotoxin MVIIA in the absence of capsaicin (Fig. 4d). Intraplantar ω-conotoxin MVIIA (0.1 μM, 1 μM, and 10 μM) did not impair behavioral responses to thermal stimuli applied 15-30 minutes after injection (n = 4-11 mice / dose; *p* = 0.835, Fig. 4d) which is evidence that ω-conotoxin MVIIA does not reach the spinal cord, at least during the time course of the behavioral measurements. Therefore, we conclude that peripheral Ca_V_2.2 channels in skin are required for normal capsaicin-induced thermal hyperalgesia.

### P2X7 receptor activation induces hyperalgesia downstream of Ca_V_2.2 channel activation

Ca_V_2.2 channels support voltage- and calcium-dependent release of transmitters from nociceptor presynaptic terminals (Figs. 2, 3) and our findings suggest that Ca_V_2.2 channels serve a similar function at nociceptor nerve endings in the skin. Several inflammatory factors are released from nociceptor free nerve endings and contribute to neurogenic inflammation. We chose to assess the role of ATP and P2X7 receptor signaling in capsaicin-induced thermal hyperalgesia based on substantial evidence indicating calcium-dependent release of ATP from neurons,^40–42^ the ability of ATP signaling to induce hyperalgesia,^43–45^ and the availability of a P2X7-specific antagonist^46,47^ and agonist.^17^ We first determined whether ATP signaling through P2X7 receptors is involved in capsaicin-induced thermal hyperalgesia using the P2X7 antagonist A438079. Thermal hyperalgesia was greatly attenuated in WT mice injected with a mixture of capsaicin and A438079 relative to capsaicin alone (WT −6.58 ± 18.5% change in response latency from baseline; 91.3 ± 24.5% decrease in effect size; *p* = 0.003, n = 9 mice, Fig 5a). These data establish a role for local P2X7 receptor activation in capsaicin-induced thermal hyperalgesia and suggest that TRPV1 channel activation triggers the release of ATP, the endogenous ligand for P2X7 receptors, from nociceptor peripheral nerve endings.

**Figure 5.**
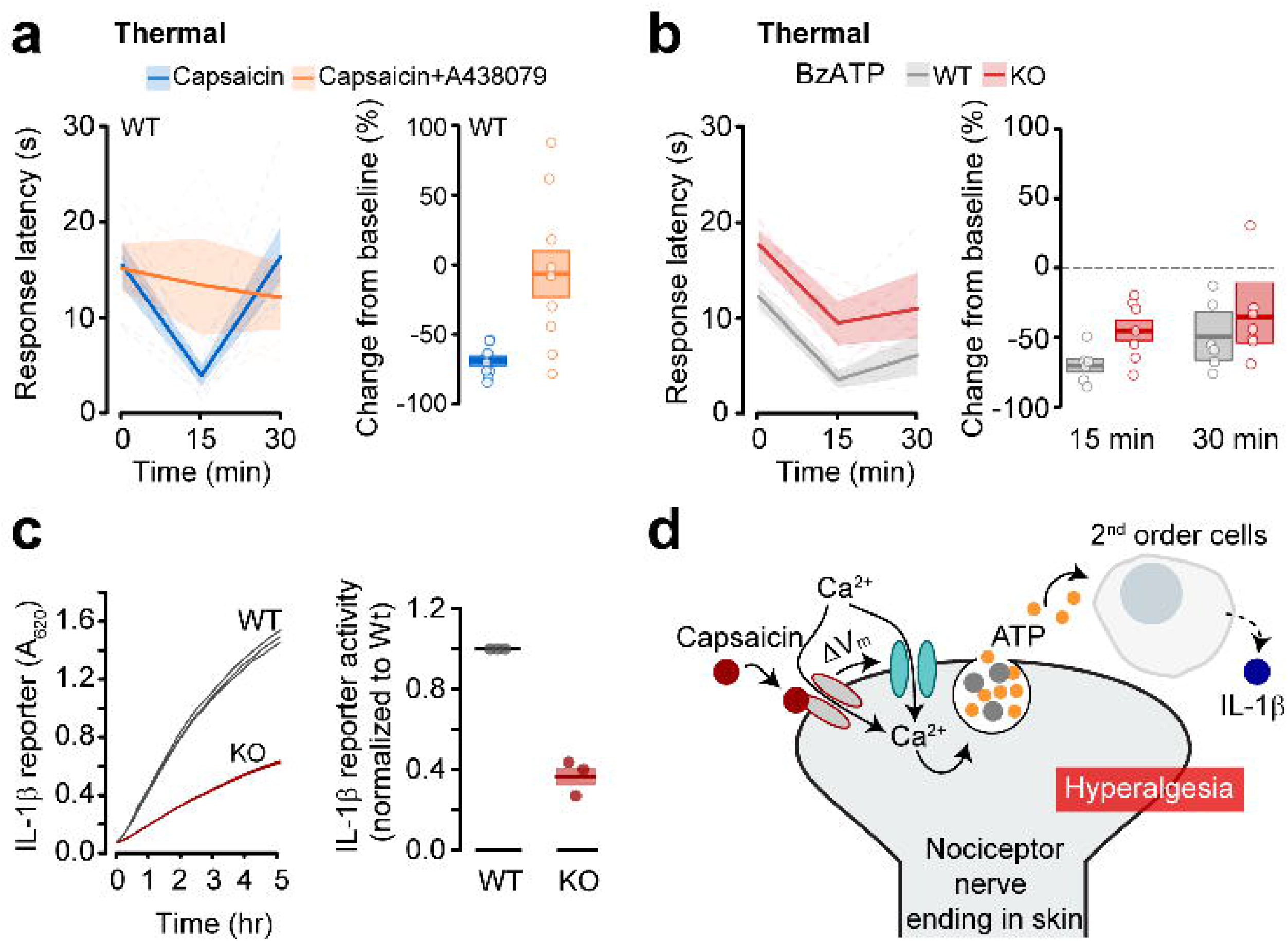
Capsaicin-induced actions of ATP depend on Ca_V_2.2 channels. **a**. Thermal sensitivity of plantar hind paw of WT mice measured before (0 min), and 15 min and 30 mins after injection of 0.1% w/v capsaicin (*blue*, n = 33 same data as in **4a**) or 0.1% w/v capsaicin + A438079 (*orange*, n = 9). All experiments were conducted under blinded conditions. *Left*, responses from individual mice (thin lines), average data (thick lines), and SE (shaded areas). *Right*, thermal responses represented as change from baseline for individual mice (symbols), average (horizontal line), and SE (shaded areas). Mean and SE were −6.58 ± 18.5%, *p* = 0.003 (t-test) compared to WT. **b**. Thermal sensitivity of plantar hind paw of WT (*gray*, n = 6) and KO (*red*, n = 7) mice measured before (0 min), and 15 min and 30 mins after injection of 5 mM BzATP. *Left*, responses from individual mice (thin lines), average data (thick lines), and SE (shaded areas). *Right*, thermal responses represented as change from baseline for individual mice (symbols), average (horizontal line), and SE (shaded areas). Mean and SE were at 15 min for WT: −70 ± 5.2% and KO: −45.0 ± 8.1%, p = 0.027 (t-test) and, at 30 min for WT: −50.6 ± 10.1% and KO: - 36.3 ± 12.2%, p = 0.388 (t-test). **c.** Continuous absorbance values for IL-1β reporter using cell-based assay, measuring release of IL-1β from plantar hind paw skin explant over a 5 hr time period. *Left* Data shows technical triplicates from WT and KO explants. *Right*, normalized values from 3 WT and 3 KO mice (symbols), average (horizontal line), and SE (shaded), *p* = 0.004 (t-test). **d**. Proposed mechanism for Ca_V_2.2 channel involvement in capsaicin-induced thermal hyperalgesia. Capsaicin binds TRPV1 channels on *Trpv1* nociceptor nerve endings in skin, the nociceptor membrane depolarizes, Ca_V_2.2 channel open, calcium enters nociceptors through Ca_V_2.2 channels triggering ATP release from secretory vesicles, ATP activates P2X7 receptors on non-neuronal, second order cell (s), and IL-1β is released. Inflammatory mediators trigger transient hyperalgesia.

In order to localize Ca_V_2.2 channels within this pathway from capsaicin to thermal hyperalgesia via P2X7 receptor activation, we used a potent P2X7 receptor agonist, BzATP, to elicit thermal hyperalgesia in WT and KO mice. BzATP induced thermal hyperalgesia in WT and KO mice (WT: −70.0 ± 5.2% change in response latency from baseline, n = 6 mice; KO: −45.0 ± 8.1% change in response latency from baseline; n = 7 mice) 15 min after injection and was sustained 30 min after injection in both strains (WT: −50.6 ± 10.1% change in response latency from baseline; n = 6 mice; KO: −36.3 ± 12.2% change in response latency from baseline; n = 7 mice). At the 15 min post-injection time point, the effect size of BzATP and capsaicin-induced thermal hyperalgesia in WT mice were not distinguishable (1.2 ± 8.8% increase in effect size; *p* = 0.999) and, in KO mice the effect size of BzATP-induced hyperalgesia was significantly greater than that induced by capsaicin (545 ± 10.5% increase in effect size; *p* = 0.014). The effect size of BzATP-mediated hyperalgesia in KO mice was similar to WT (35.7 ± 14.0% decrease KO effect size relative WT effect size; *p* = 0.543). These data suggest that P2X7 receptor activation triggers hyperalgesia via a pathway that is downstream of and independent of Ca_V_2.2 channel activation.

Our results show that capsaicin-induced hyperalgesia depends on activation of both local Ca_V_2.2 channels and local P2X7 receptors, and that ATP acts via P2X7 receptors primarily downstream of Ca_V_2.2 channel activation.

A role for P2X7 receptors in capsaicin-evoked hyperalgesia implicates non-neuronal cells, because P2X7 receptors are primarily expressed by immune cells. P2X7 receptor activation triggers the release of the prominent inflammatory mediator IL-1β.^48^ We therefore measured levels of IL-1β released from isolated skin samples from WT and KO mice over 72 hr. IL-1β levels in conditioned media from skin of KO mice were 40% of WT (Fig 5c), suggesting that Ca_V_2.2 channels in skin may also contribute to inflammatory hyperalgesia via regulation of immune cell activity (Fig 5d).

In summary, our data show that Ca_V_2.2 channels in nociceptor termini in skin are critical, early in the neurogenic inflammatory response pathway, for locally induced thermal hyperalgesia. This new role expands the importance of Ca_V_2.2 channels in signal transmission in nociceptors beyond its well documented role in supporting transmitter release at pre-synaptic terminals. Furthermore, we show that the Ca_V_2.2 channel splice isoform containing e37a may have a unique role in thermal hyperalgesia resulting from capsaicin-mediated neuronal activation that is not fully supported by Ca_V_2.2 e37b splice isoforms.

## DISCUSSION

Ca_V_2.2 channels are important targets of analgesic drugs and neuromodulators that regulate calcium entry at presynaptic terminals.^1,12,49,50^ In this study, we reveal a previously undocumented role of Ca_V_2.2 channels filling a knowledge gap between the first step in neurogenic inflammatory signaling and thermal hyperalgesia. We show that peripheral Ca_V_2.2 channels in skin are required for short-term capsaicin-induced thermal hyperalgesia, but not for mechanical hyperalgesia. Our data are consistent with a model (Fig. 5d) in which Ca_V_2.2 channels open in response to nociceptor membrane depolarization and, following capsaicin exposure, proinflammatory factors, including ATP, are released. ATP acts via P2X7 receptors of non-neuronal cells to induce the release of inflammatory mediators including secreted IL-1β. Our data are consistent with the analgesic actions of locally applied combined inhibitors of Ca_V_2.2 and Na_V_1.7 channels in different mouse models of inflammatory pain.^24^

While previous studies have focused almost exclusively on centrally located Ca_V_2.2 channels, we show that peripheral Ca_V_2.2 channels in skin are essential for thermal hyperalgesia, a finding that may help explain previous findings.^32–34^ For example, studies of KO mice show reduced hyperalgesia induced by formalin but in some of these studies, this occurs independent of effects on acute nociception.^32–34^ Our data resolve these apparent inconsistent observations, by showing that peripheral Ca_V_2.2 channels support hyperalgesia, independent of their role in acute basal nociception at central sites.

### Peripheral Ca_V_2.2 channels in skin required for thermal hyperalgesia

Capsaicin-induced hyperalgesia is a robust model of transient neurogenic inflammation. However, little or no attention has been paid to understanding how capsaicin-induced depolarization of *Trvp1*-nociceptors triggers an inflammatory response.^51^ Ca_V_2.2 channels in peripheral nerve termini are activated by capsaicin-induced depolarization, either directly and/or by back propagating action potentials.^15,52^ Exocytosis of peptidergic and ATP-containing vesicles is widely acknowledged to follow nociceptor activation^16,23,53^ and our study demonstrates that Ca_V_2.2 channels are the critical link between these two steps in neurogenic thermal hyperalgesia.^18,54,55^

Local block of Ca_V_2.2 channels in skin by ω-conotoxin MVIIA significantly, and specifically attenuated capsaicin-induced thermal hyperalgesia in mice while leaving mechanical hyperalgesia and basal nociception intact. This is consistent with the divergent signaling pathways that support thermal and mechanical hyperalgesia. ^56–61^Interestingly, a combined inhibitor of Ca_V_2.2 and Na_V_1.7 channels applied at peripheral sites in animal models of more prolonged inflammation were found to be effective on both thermal and mechanical hyperalgesia.^24^ It will be interesting to tease out the differences in the role of Ca_V_2.2 channels in the induction of transient and maintenance of more prolonged forms of hyperalgesia.

Consistent with these data, we also show that the Ca_V_2.2-e37a channel splice isoform may have unique properties that are important in capsaicin-induced thermal hyperalgesia, which are not complexly substituted for by Ca_V_2.2-e37b channel isoforms.

Ca_V_2.2 channels couple membrane depolarization to exocytosis of a range of neurotransmitter containing vesicles^5,13,40,41,49,62,63^ and we present evidence that Ca_V_2.2 channels in *Trpv1*-nociceptor nerve endings in skin trigger ATP release to support capsaicin-induced thermal hyperalgesia (Fig 5).^18,19,42,64^ We show that P2X7 receptor blockade inhibits the response in WT mice,^65,66^ while intraplantar BzATP mimics the response in KO mice; the latter observation is consistent with a role for Ca_V_2.2 channel activation in capsaicin-induced thermal hyperalgesia upstream of ATP release.

ATP is packaged into secretory vesicles and acts as a neurotransmitter to influence the activity of nociceptors, glia, and immune cells.^40^ P2X7 receptors are located on non-neuronal cells, thus our data also implicate non-neuronal cells in the development of transient thermal hyperalgesia induced by intraplantar capsaicin (Fig. 5). Although capsaicin-induced thermal hyperalgesia was eliminated by local P2X7 receptor blockage, our results do not exclude the involvement of other inflammatory signals or signaling pathways. Inflammatory molecules, in addition to ATP, such as neuropeptides can act in synergy or downstream of ATP^23^ and Ca_V_2.2 channels could contribute to ongoing release of factors from nociceptor nerve endings or microglia.^67^ During prolonged inflammation, ATP can also be released from lysosomes, dead and dying cells, and diffusion through large hemi-channel pores.^18,19,42,64^

### Peripheral inhibition of Ca_V_2.2 channels for analgesia

Ca_V_2.2 channels are important targets for the development of novel analgesics as evidenced by the initial clinical success of ω-conotoxin MVIIA as an analgesic in patients with otherwise intractable chronic pain.^3–6,68^ Our data raise the possibility that local injection of ω-conotoxin MVIIA, or, as shown by others, dual inhibitors of Ca_V_2.2 and Na_V_1.7,^24^ may have clinical utility against certain inflammatory responses, by interrupting signals locally, thereby avoiding severe, debilitating side-effects^3,6,9,69^ that limit that current use of Ca_V_2.2 channel blockers.^70^

Intraplantar ω-conotoxin MVIIA was highly effective at inhibiting thermal hyperalgesia independent of effects on basal nociception, central Ca_V_2.2 channels (Fig. 4). Local injections of anesthetics, steroids, and botulinum toxin are used routinely to manage a range of pain conditions.^71^ Although long-term safety of locally applied ω-conotoxin MVIIA in various tissue in patients would need to be tested, published clinical data suggests a high likelihood of tolerability and efficacy.^4,70,72^

### Peripheral Ca_V_2.2 channel slice isoforms

We also demonstrate that Ca_V_2.2-e37a channel splice isoforms expressed in *Trvp1*-nociceptors have properties that are not fully substituted for by Ca_V_2.2-e37b channels, with respect to their role in capsaicin-induced thermal hyperalgesia (Fig. 4). By contrast, Ca_V_2.2-e37a and Ca_V_2.2-e37b channels appear to be fully interchangeable with respect to their function in supporting acute nociception (Fig. 1–3).^14^ Compared to Ca_V_2.2-e37b, Ca_V_2.2-e37a channels are trafficked more efficiently to the cell surface, and they are more susceptible to Gαi/o dependent inhibition including by μ-opioid receptors.^14,28,29,73^ We speculate, but it remains to be tested, that Ca_V_2.2-e37a isoforms are trafficked with greater efficiency to peripheral nerve endings in skin, as compared to Ca_V_2.2-e37b isoforms. Nonetheless, based on our data, isoform-selective inhibitors specifically targeting Ca_V_2.2-e37a channels might be expected to preferentially act on nociceptor Ca_V_2.2 channels, with few off-target effects and relatively less impact on normal, _acute nociception._^14,27,28,31^

Ca_V_2.2-e37a channels are enriched in *Trpv1*-expressing nociceptors and *Cacna1b*-e37a mRNAs are at low or undetectable levels elsewhere in the nervous system.^27^ We recently elucidated the molecular mechanism controlling Ca_V_2.2-e37a channel expression in *Trpv1*-nociceptors. Cell-specific hypomethylation of e37a locus in *Cacna1b*, allows CTCF to bind, promoting e37a recognition and its inclusion in *Cacna1b* mRNA in *Trpv1*-nociceptors.^30^ By contrast, in non *Trpv1* cell population and in *Trpv1*-expressing nociceptors 7 days after nerve injury, e37a locus is methylated, CTCF does not bind, and e37a is skipped. It will be interesting to know whether chronic inflammatory models similarly disrupt *Cacna1b* e37a splicing.

Much progress has been made in identifying and inhibiting, the downstream actions of inflammatory mediators that are released in response to chemical irritants, tissue injury, and certain diseases. Our studies show that Ca_V_2.2 channels are critical for the action of early inflammatory mediators ATP and IL-1β, both of which are implicated in several animal models of hyperalgesia. Inflammation is a ubiquitous response to tissue damage and a significant contributor to chronic, pathological pain in human patients. A more precise understanding of peripheral Ca_V_2.2 channels advances our basic understanding of factors that contribute to inflammatory pain and may suggest improved therapies for certain pathological pain conditions.

## MATERIALS AND METHODS

All mice used in this work were bred at Brown University and all protocols and procedures were approved by the Brown University Institutional Animal Care and Use Committee. For all behavioral experiments, both male and female mice were included. For physiology experiments, sex was not identified prior to tissue collection. All values shown are mean ± standard error. All behavior experiments assessing mechanical sensitivity were conducted under blinded conditions and, where indicated, thermal sensitivity experiments were conducted under both blinded and non-blinded conditions by independent experimenters in distinct cohorts of mice. Although large differences in general activity level between Ca_V_2.2-null (KO) and WT mice made genotype blinding ineffective, in blinded experiments, experimenter was blinded to both mouse genotype and/or injection.

### Mice

*Trpv1-Cre*^74^ (RRID: IMSR_JAX:017769), *lox-STOP-lox-ChR2-EYFP*^75^ (RRID: IMSR_JAX:012569, hereafter *ChR2-EYFP*), and *lox-STOP-lox-tdTomato*^76^ (RRID: IMSR_JAX:007908, hereafter *tdTomato*) mice were purchased from The Jackson Laboratory. *Trpv1* is expressed in testes during gamete production,^77–79^ inducing Cre-dependent reporter expression in spermatozoa. We confirmed widespread reporter expression beyond the *Trpv1*-lineage in offspring of *Trpv1*^Chr2EYFP+/-^ or *Trpv1*^tdTomato+/-^ mice; therefore, all mice used in this study were first-generation progeny of single homozygous parents. *Trpv1*-Cre^+/+^ mice were mated with either *ChR2-EYFP*^+/+^ or *tdTomato^+/+^* mice to generate *Trpv1*^Chr2EYFP+/-^ or *Trpv1*^tdTomato+/-^ offspring. Generation of *Cacna1b*^37b-37b^ mice was described previously.^14^

We generated a Ca_V_2.2-null mouse strain by insertion of an EGFP+stop cassette in frame in exon 1 of the *Cacna1b* gene. All primers used in creation of Ca_V_2.2-null mouse strain are shown in Table 1. To create the Cacna1b^-/-^ targeting construct, a 12 kb NsiI fragment from the 129S mouse BAC genomic clone bMQ122-B9 (Source BioScience) was cloned into the PstI site of pBSSK^+^. An AgeI site was inserted into exon 1 of *Cacna1b* by mutagenesis using a QuickChange II XL site-directed mutagenesis kit (Stratagene) with primers 1-For and 1-Rev. The EGFP+stop cassette was inserted in-frame into exon 1 at the AgeI site following amplification from pEGFP-C1 (BD Biosciences) using primers 2-For and 2-Rev. The loxP-NeoR-loxP cassette was inserted at the MluI site in the intron between exons 1 and 2 following PCR amplification from pL452 (Addgene) using primers 3-For and 3-Rev. The final targeting construct was 14.8kb.

**Table 1:**
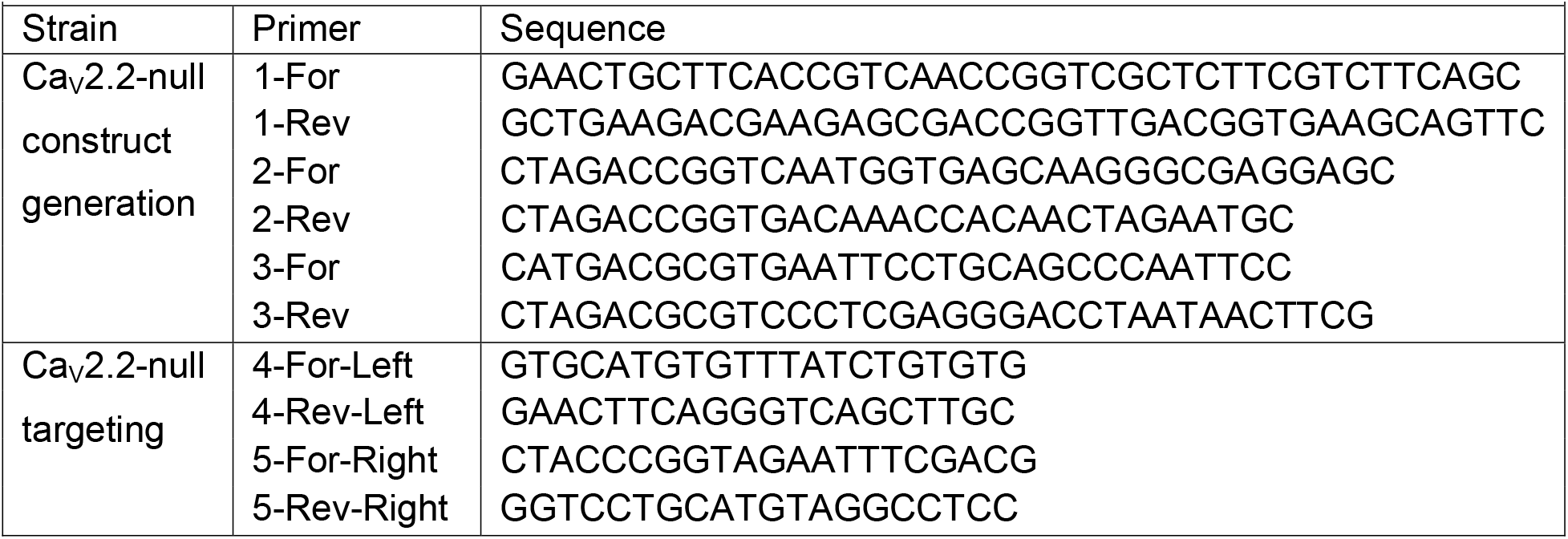
Primers for generation of Ca_V_2.2-null mouse strain.

Mouse 129Ola ES cells derived from a male embryo were grown on mitotically inactive SNL76/7 feeder cells. Ten million (10^7^) ES cells were electroporated with 20 μg of a construct linearized with PvuI, and G418 selection was initiated after 24 h. Correctly targeted ES cell clones were identified by PCR and injected in E3.5 blastocysts isolated from C57Bl/6-*Tyrc-Brd* female mice. Injected blastocysts were implanted into day 2.5 pseudopregnant females for the generation of chimeras. Male chimeras were mated with *C57BL/6-Tyrc-Brd* females to obtain F1 progeny. Germ line transmission was confirmed by PCR of genomic DNA from each ES cell clone to confirm homologous integration from the long arm and short arm of the targeting vector using primers indicated in Table 1 for the left and right arms. The neomycin resistance cassette was subsequently removed from F1 mice by crossing to a B6.FVB-Tg(EIIa-cre)C5379Lmgd/J (RRID: IMSR_JAX:003724). Deletion of the neomycin cassette was confirmed by PCR amplification of genomic DNA.

Individual WT strains were generated in parallel from the same genetic background used to create *Cacna1b^-/-^* and *Cacna1b^37b/37b^* mouse strains. The two WT strains were pooled together following determination that there were no significant differences on any assessments.

### Acute nociception assays

Responses to thermal, mechanical, and LED stimulation were analyzed from male and female *Cacna1b*^+/+^/*Trpv1*^Chr2EYFP+/-^, *Cacna1b*^-/-^/*Trpv1*^Chr2EYFP+/-^, and *Cacna1b*^37b-37b^/*Trpv1*^Chr2EYFP+/-^. Mice were between 2 and 4 months old. In total, 195 *Cacna1b*^+/+^/*Trpv1*^Chr2EYFP+/-^ mice, 89 *Cacna1b^-/-^* /*Trpv1*^Chr2EYFP+/-^, and 75 *Cacna1b*^37b-37b^/*Trpv1*^Chr2EYFP+/-^ mice were used to monitor acute nocifensive responses: thermal (146 WT, 56 Ca_V_2.2-null, 42 Ca_V_2.2-e37b), mechanical (32 WT, 24 Ca_V_2.2-null, 26 Ca_V_2.2-e37b), LED (17 WT, 9 Ca_V_2.2-null, 7 Ca_V_2.2-e37b).

We used a Plantar Analgesia Meter (IITC) to assess thermal responses to radiant heat. Mice were placed in Plexiglas containers on an elevated glass plate and allowed to habituate for 1 hr prior to testing. A visible-light, radiant heat source was positioned beneath the mice and aimed using low-intensity visible light to the plantar surface of the hindpaw. An orange-pass filter was used to prevent blue-light activation of channelrhodopsin in sensory nerve terminals. Trials began once the high-intensity light source was activated and ended once the mouse withdrew their hindpaw and 1) shook their paw, 2) licked their paw, or 3) continued to withdraw their paw from stimulation. Immediately upon meeting response criteria, the high-intensity light-source was turned off. The latency to response was measured to the nearest 0.01 s for each trial using the built-in timer, which is activated and de-activated with the high-intensity beam. For all trials, high-intensity beam was set at 40%, low-intensity beam set at 10%, and maximum trial duration was 30s. Three trials were conducted on each hindpaw for each mouse, with at least 1 minute between trials of the same hindpaw.^80^

To assess responses to mechanical stimuli, we used a Dynamic Plantar Aesthesiometer (Ugo Basile, Cat# 37450). Mice were placed in a Plexiglas container over an elevated mesh platform at least 30 minutes to allow accommodation before the measurement. The plantar surface of the hind paw was stimulated with a steel rod pushed against the plantar side of the hind paw with linear ascending force, from 0 to 5 g over 10 s, in 0.5 g/s intervals, until a fast withdraw of the paw was observed or reach a cut-off time of 20 s. The latency to response in seconds and actual force to the nearest 0.1 g at the time of paw withdrawal are automatically detected and recorded by the unit. Three trials were conducted on each hind paw for each mouse, with at least 1 minute between trials of the same hind paw.

For assessing nocifensive responses to optogenetic nociceptor activation, male or female mice were placed in Plexiglas containers on an elevated glass surface. A fiber-coupled, blue (465 nm) LED light (Plexon) was mounted to a movable stage at a fixed distance below the glass platform. Light intensity was controlled by the supplied driver and intensity (1-9 mW, ~0.1-1 mW/mm^2^) was measured using a light meter (PM100A, Sensor: S121C, Thor Labs) mounted on top of the glass plate. Mice were allowed to habituate to the chamber for 1 hr prior to testing. Each mouse was stimulated 10 times, equally divided between the left and right hindpaw, at each of 6 light intensity levels, for a total of 60 trials per mouse. In each trial, the LED light was directed at the plantar surface of one hindpaw for 5 seconds or until a nocifensive response was elicited. Nocifensive responses included hindpaw withdrawal accompanied by at least one of the following: shaking or licking the stimulated hindpaw or continued hindpaw withdrawal during prolonged stimulation. All responses occurred within 3 seconds of stimulus onset. The number of nocifensive trials was counted for each mouse at each light intensity level and response distributions for individual mice were fit using a 4-parameter logistic curve:

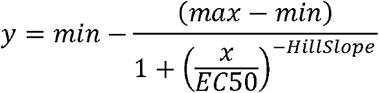

Values for minimum, maximum, EC50, and HillSlope were calculated automatically and mean values were used to construct a fit of the average response distribution.

### Immunohistochemistry

*Cacna1b*^+/+^/*Trpv1*^Chr2EYFP+/-^ and *Cacna1b*^+/+^/*Trpv1*^tdTomato+/-^ male mice (2-6 months) were transcardially perfused with cold phosphate buffered saline (PBS), followed by 4% paraformaldehyde (PFA) in PBS. Spinal cord (L4-L6) and skin (hindpaw) were removed and post-fixed in 4% PFA overnight at 4°C. Samples were cryoprotected in 30% sucrose in PBT at 4°C for 48 hrs, frozen in OCT media, and cut into 14 μm slices.

Slices were blocked overnight using 5% bovine serum albumin (BSA, Sigma Aldrich) in PBS with 0.4% Triton X-100 (Sigma, PBST). anti-CGRP (Millipore Cat# PC205L, RRID: AB_2068524) antibody was applied in PBST with 5% BSA at 1:250 dilution for 48 hrs at 4°C. Secondary antibodies (Alexa488 donkey-anti-rabbit (RRID: AB_2571722), Alexa488 donkey-anti-mouse (RRID: AB_2571721)) were applied at 1:200 in PBST with Alexa647-conjugated Isolectin B4 (SCR_014365) at 1:100 and DAPI (SCR_014366) at 1:1000 for 4 hr at room temperature. Images were collected using a Zeiss LSM 800 confocal microscope using ZEN software.

### RNAScope in situ hybridization

RNAScope (^®^, Advanced Cell Diagnostics) in situ hybridization was performed as instructed in the manual, with the following differences. DRG were isolated from *Cacna1b*^+/+^ and *Cacna1b*^-/-^ and *Cacna1b*^37b-37b^ mice, and processed and sectioned as described above (Methods/Immunohistochemistry). Following sectioning, slices were dehydrated with 100% ethanol for 5 min and allowed to air dry. A barrier was drawn around each section using a hydrophobic barrier pen. Slides were incubated at 40°C in Protease III for 30 min. *Cacna1b* or *Cacna1b*-e37a probes were applied at 40°C for 4 hrs and detected using amplification reagents 1-4, as described in the kit. Images were collected using a Zeiss LSM 800 confocal microscope using ZEN software. At least 3 slides from 2 mice (2 months old) were used for analysis. Probe specificity was confirmed using Ca_V_2.2-null and Ca_V_2.2-e37b samples.

### Spinal cord slice physiology

Acute spinal cord slices were prepared from *Cacna1b*^+/+^/*Trpv1*^Chr2EYFP+/-^, *Cacna1b^-/-^*/*Trpv1*^Chr2EYFP+/-^, and *Cacna1b*^37b-37b^/*Tpv1*^Chr2EYFP+/-^ mice (mixed gender, p14-21). Mice were anesthetized by ip injection of Beuthanasia-S and transcardially perfused with cold, oxygenated aCSF containing, in mM: 125 NaCl, 2.5 KCl, 26 NaHCO3, 1.25 NaH2PO4, 1.5 CaCl2, 6 MgCl2, 25 Glucose, 1 Kynurenic acid. The spinal cord was removed, embedded in 2% low-melting agarose, and cut into 300 um sections using a Leica VT1200S vibrating blade microtome in the same aCSF solution. Sections were then transferred to holding chamber containing the same aCSF solution at 30°C for 1 hr and at room temperature thereafter. For recording, individual slices were transferred to a recording chamber and continually perfused with oxygenated aCSF containing, in mM: 119 NaCl, 2.5 KCl, 26 NaHCO3, 1 NaH2PO4, 2.5 CaCl2, 1.3 MgSO4, 25 Glucose, 1.3 Na-Ascorbate. Patch pipettes were filled with an internal solution containing, in mM: 125 KGluconate, 28 NaCl, 2 MgCl2, 2 Mg-ATP, 0.3 Na-GTP, 0.6 EGTA, 10 HEPES and had a resistance between 3-5 MΩ. Synaptic responses were elicited by 1 ms pulses of blue light through the 40x microscope objective controlled by a shutter and the blue light was focused directly on the recorded cell. Cells were held at −70 mV during all protocols.

### Inflammatory models

*Cacna1b*^+/+^/*Trpv1*^Chr2EYFP+/-^, *Cacna1b*^-/-^/*Trpv1*^Chr2EYFP+/-^, and *Cacna1b*^37b-37b^/*Trpv1*^Chr2EYFP+/-^ mice (male and female, 4-6 months) were anesthetized with isoflurane. Capsaicin (0.1% w/v, in sterile saline with 5% Tween 20, 20 μL), BzATP (5 mM in water, 20 μL), A438079 (6 mM in sterile saline, 20 μL), or ω-conotoxin MVIIA (1 μM in sterile saline, 20 μL) was injected via 30-gauge needle into the left hindpaw and mice were allowed to recover in the testing chamber. Thermal or mechanical sensitivity was assessed using the radiant heat or automated von Frey assays, respectively (described above). Median response latency of at least three trials at each time point was used to assess sensitivity. To compared effect size between different mouse strains we applied the following formula: ((Effect(Mouse A) - Effect(WT)) / Effect(WT)) x 100%.

### IL-1β analysis

*Cacna1b*^+/+^/*Trpv1*^Chr2EYFP+/-^ and *Cacna1b*^-/-^*Trpv1*^Chr2EYFP+/-^ mice (n = 3 mice per genotype) were anesthetized with isoflurane and euthanized via i.p. injection of euthanasia solution. The plantar skin from the left and right hindpaws was removed and ~20 mg of skin was placed in 1 mL DMEM supplemented with 10% heat-inactivated fetal bovine serum and Penicillin-Streptomycin (Gibco). Skin cultures were maintained at 37°C with 5% CO_2_ for 72 hrs. One day prior to collection of conditioned media, HEK-Blue IL-1β cells (InvivoGen) were split into a 24-well plate with 0.5 mL DMEM supplemented with 10% heat-inactivated fetal bovine serum, Hygromycin, and Zeocin. Conditioned media containing IL-1β was collected from skin cultures and 200 μL was added to each well of HEK-Blue cells for 24 hrs. Secreted embryonic alkaline phosphatase was detected by mixing 200 μL Quanti-Blue media with 50 μL HEK-Blue cell media in triplicate in a 96-well plate and quantified by reading the absorbance at 620 nm using a Biotek Synergy HTX multi-mode microplate reader at 37°C. Quanti-Blue media alone was used as a blank sample.

### Statistical analyses

Figure 1c, 4, 5a-b: Statistical significance of percent change from baseline at 15 minutes post injection was assessed in R (v3.5.0)/RStudio (v1.1.463) using univariate ANOVA (type III sums of squares) with Tukey HSD post-hoc correction for multiple comparisons. When both blinded and non-blinded experiments were conducted, a significant effect of cohort was first established using univariate ANOVA with type III sums of squares and cohorts were pooled if no main effect or interaction involving cohort was observed.

Figure 1d, 2, 3: Analyses were performed using IBM SPSS Statistics 24. Repeated measures ANOVA was used to assess sensitivity to optogenetic stimulation. Synaptic current properties were assessed together using Multivariate ANOVA to correct for correlation between measured parameters from the same recording. For correction of multiple comparisons between genotypes, Dunnett T3 and Tukey HSD were used. Dunnett T3 was used when the homogeneity of group variances assumption was not met.

## Acknowledgements

The authors thank Sylvia Denome for expert technical assistance. This work was funded by grants NS055251 (DL), F31NS093818 (DD) and Warren Alpert Fellowship Award and K99NS116123 (EJLS).

## Notes

### Competing Interest Statement

The authors have declared no competing interest.

## REFERENCES

1. Motin, L. & Adams, D. J. omega-Conotoxin inhibition of excitatory synaptic transmission evoked by dorsal root stimulation in rat superficial dorsal horn. Neuropharmacology 55, 860–864 (2008).

2. Patel, R., Montagut-Bordas, C. & Dickenson, A. H. Calcium channel modulation as a target in chronic pain control: Calcium channel antagonists and chronic pain. Br. J. Pharmacol. (2017) doi:10.1111/bph.13789.

3. Yekkirala, A. S., Roberson, D. P., Bean, B. P. & Woolf, C. J. Breaking barriers to novel analgesic drug development. Nat. Rev. Drug Discov. (2017) doi:10.1038/nrd.2017.87.

4. McGivern, J. G. Ziconotide: a review of its pharmacology and use in the treatment of pain. Neuropsychiatr. Dis. Treat. 3, 69–85 (2007).

5. Zamponi, G. W., Striessnig, J., Koschak, A. & Dolphin, A. C. The Physiology, Pathology, and Pharmacology of Voltage-Gated Calcium Channels and Their Future Therapeutic Potential. Pharmacol. Rev. 67, 821–870 (2015).

6. Lee, S. Pharmacological Inhibition of Voltage-gated Ca(2+) Channels for Chronic Pain Relief. Curr. Neuropharmacol. 11, 606–620 (2013).

7. Bowersox, S. S. et al. Selective N-type neuronal voltage-sensitive calcium channel blocker, SNX-111, produces spinal antinociception in rat models of acute, persistent and neuropathic pain. J. Pharmacol. Exp. Ther. 279, 1243–1249 (1996).

8. Wang, Y. X., Pettus, M., Gao, D., Phillips, C. & Scott Bowersox, S. Effects of intrathecal administration of ziconotide, a selective neuronal N-type calcium channel blocker, on mechanical allodynia and heat hyperalgesia in a rat model of postoperative pain. Pain 84, 151–158 (2000).

9. Bannister, K., Kucharczyk, M. & Dickenson, A. H. Hopes for the Future of Pain Control. Pain Ther. (2017) doi:10.1007/s40122-017-0073-6.

10. Barzan, R., Pfeiffer, F. & Kukley, M. N- and L-Type Voltage-Gated Calcium Channels Mediate Fast Calcium Transients in Axonal Shafts of Mouse Peripheral Nerve. Front. Cell. Neurosci. 10, (2016).

11. Regan, L. J., Sah, D. W. & Bean, B. P. Ca2+ channels in rat central and peripheral neurons: high-threshold current resistant to dihydropyridine blockers and omega-conotoxin. Neuron 6, 269–280 (1991).

12. Kress, M., Izydorczyk, I. & Kuhn, A. N-and L-but not P/Q-type calcium channels contribute to neuropeptide release from rat skin in vitro. Neuroreport 12, 867–870 (2001).

13. Kamp, M. A., Hänggi, D., Steiger, H.-J. & Schneider, T. Diversity of presynaptic calcium channels displaying different synaptic properties. Rev. Neurosci. 23, 179–190 (2012).

14. Andrade, A., Denome, S., Jiang, Y.-Q., Marangoudakis, S. & Lipscombe, D. Opioid inhibition of N-type Ca2+ channels and spinal analgesia couple to alternative splicing. Nat. Neurosci. 13, 1249–1256 (2010).

15. Blair, N. T. & Bean, B. P. Roles of tetrodotoxin (TTX)-sensitive Na+ current, TTX-resistant Na+ current, and Ca2+ current in the action potentials of nociceptive sensory neurons. J. Neurosci. Off. J. Soc. Neurosci. 22, 10277–10290 (2002).

16. Louis, S. M., Jamieson, A., Russell, N. J. & Dockray, G. J. The role of substance P and calcitonin gene-related peptide in neurogenic plasma extravasation and vasodilatation in the rat. Neuroscience 32, 581–586 (1989).

17. Jarvis, M. F. et al. Modulation of BzATP and formalin induced nociception: attenuation by the P2X receptor antagonist, TNP-ATP and enhancement by the P2X _3_ allosteric modulator, cibacron blue. Br. J. Pharmacol. 132, 259–269 (2001).

18. Jung, J., Jo, H. W., Kwon, H. & Jeong, N. Y. ATP release through lysosomal exocytosis from peripheral nerves: the effect of lysosomal exocytosis on peripheral nerve degeneration and regeneration after nerve injury. BioMed Res. Int. 2014, 936891 (2014).

19. Cook, S. P. & McCleskey, E. W. Cell damage excites nociceptors through release of cytosolic ATP. Pain 95, 41–47 (2002).

20. Koltzenburg, M., Bennett, D. L., Shelton, D. L. & McMahon, S. B. Neutralization of endogenous NGF prevents the sensitization of nociceptors supplying inflamed skin. Eur. J. Neurosci. 11, 1698–1704 (1999).

21. Woolf, C. J., Safieh-Garabedian, B., Ma, Q. P., Crilly, P. & Winter, J. Nerve growth factor contributes to the generation of inflammatory sensory hypersensitivity. Neuroscience 62, 327–331 (1994).

22. Ma, Q. P. & Woolf, C. J. The progressive tactile hyperalgesia induced by peripheral inflammation is nerve growth factor dependent. Neuroreport 8, 807–810 (1997).

23. Chiu, I. M., von Hehn, C. A. & Woolf, C. J. Neurogenic inflammation and the peripheral nervous system in host defense and immunopathology. Nat. Neurosci. 15, 1063–1067 (2012).

24. Lee, S. et al. Novel charged sodium and calcium channel inhibitor active against neurogenic inflammation. eLife 8, (2019).

25. Gray, A., Raingo, J. & Lipscombe, D. Neuronal calcium channels: Splicing for optimal performance. Cell Calcium 42, 409–417 (2007).

26. Lipscombe, D. Neuronal proteins custom designed by alternative splicing. Curr. Opin. Neurobiol. 15, 358–363 (2005).

27. Bell, T. J., Thaler, C., Castiglioni, A. J., Helton, T. D. & Lipscombe, D. Cell-specific alternative splicing increases calcium channel current density in the pain pathway. Neuron 41, 127–138 (2004).

28. Jiang, Y.-Q., Andrade, A. & Lipscombe, D. Spinal morphine but not ziconotide or gabapentin analgesia is affected by alternative splicing of voltage-gated calcium channel Ca_V_2.2 pre-mRNA. Mol. Pain 9, 67 (2013).

29. Macabuag, N. & Dolphin, A. C. Alternative Splicing in Ca(V)2.2 Regulates Neuronal Trafficking via Adaptor Protein Complex-1 Adaptor Protein Motifs. J. Neurosci. Off. J. Soc. Neurosci. 35, 14636–14652 (2015).

30. Lopez Soto, E. J. & Lipscombe, D. Cell-specific exon methylation and CTCF binding in neurons regulates calcium ion channel splicing and function. eLife 2020;9:e54879.

31. Altier, C. et al. Differential Role of N-Type Calcium Channel Splice Isoforms in Pain. J. Neurosci. 27, 6363–6373 (2007).

32. Hatakeyama, S. et al. Differential nociceptive responses in mice lacking the alpha(1B) subunit of N-type Ca(2+) channels. Neuroreport 12, 2423–2427 (2001).

33. Saegusa, H. et al. Suppression of inflammatory and neuropathic pain symptoms in mice lacking the N-type Ca2+ channel. EMBO J. 20, 2349–2356 (2001).

34. Kim, C. et al. Altered nociceptive response in mice deficient in the alpha(1B) subunit of the voltage-dependent calcium channel. Mol. Cell. Neurosci. 18, 235–245 (2001).

35. Debanne, D., Guérineau, N. C., Gähwiler, B. H. & Thompson, S. M. Paired-pulse facilitation and depression at unitary synapses in rat hippocampus: quantal fluctuation affects subsequent release. J. Physiol. 491 (Pt 1), 163–176 (1996).

36. Dobrunz, L. E. & Stevens, C. F. Heterogeneity of Release Probability, Facilitation, and Depletion at Central Synapses. Neuron 18, 995–1008 (1997).

37. Manabe, T., Wyllie, D. J., Perkel, D. J. & Nicoll, R. A. Modulation of synaptic transmission and long-term potentiation: effects on paired pulse facilitation and EPSC variance in the CA1 region of the hippocampus. J. Neurophysiol. 70, 1451–1459 (1993).

38. Gilchrist, H. D., Allard, B. L. & Simone, D. A. Enhanced withdrawal responses to heat and mechanical stimuli following intraplantar injection of capsaicin in rats. Pain 67, 179–188 (1996).

39. Caterina, M. J. et al. Impaired nociception and pain sensation in mice lacking the capsaicin receptor. Science 288, 306–313 (2000).

40. Pankratov, Y., Lalo, U., Verkhratsky, A. & North, R. A. Vesicular release of ATP at central synapses. Pflüg. Arch. - Eur. J. Physiol. 452, 589–597 (2006).

41. Chai, Z. et al. Ca_V_2.2 Gates Calcium-Independent but Voltage-Dependent Secretion in Mammalian Sensory Neurons. Neuron (2017) doi:10.1016/j.neuron.2017.10.028.

42. Burnstock, G. Physiology and pathophysiology of purinergic neurotransmission. Physiol. Rev. 87, 659–797 (2007).

43. Baraldi, P. G., Di Virgilio, F. & Romagnoli, R. Agonists and antagonists acting at P2X7 receptor. Curr. Top. Med. Chem. 4, 1707–1717 (2004).

44. Teixeira, J. M., Oliveira, M. C. G., Parada, C. A. & Tambeli, C. H. Peripheral mechanisms underlying the essential role of P2X7 receptors in the development of inflammatory hyperalgesia. Eur. J. Pharmacol. 644, 55–60 (2010).

45. Teixeira, J. M., de Oliveira-Fusaro, M. C. G., Parada, C. A. & Tambeli, C. H. Peripheral P2X7 receptor-induced mechanical hyperalgesia is mediated by bradykinin. Neuroscience 277, 163–173 (2014).

46. McGaraughty, S. et al. P2X7-related modulation of pathological nociception in rats. Neuroscience 146, 1817–1828 (2007).

47. Chessell, I. P. et al. Disruption of the P2X7 purinoceptor gene abolishes chronic inflammatory and neuropathic pain: Pain 114, 386–396 (2005).

48. Giuliani, A. L., Sarti, A. C., Falzoni, S. & Di Virgilio, F. The P2X7 Receptor-Interleukin-1 Liaison. Front. Pharmacol. 8, (2017).

49. Catterall, W. A. Interactions of presynaptic Ca2+ channels and snare proteins in neurotransmitter release. Ann. N. Y. Acad. Sci. 868, 144–159 (1999).

50. Yaksh, T. L. Calcium Channels As Therapeutic Targets in Neuropathic Pain. J. Pain 7, S13–S30 (2006).

51. Sweitzer, S. M. et al. Peripheral and central p38 MAPK mediates capsaicin-induced hyperalgesia. Pain 111, 278–285 (2004).

52. Castiglioni, A. J., Raingo, J. & Lipscombe, D. Alternative splicing in the C-terminus of Ca_V_2.2 controls expression and gating of N-type calcium channels: C-terminus splicing in N-type calcium channels. J. Physiol. 576, 119–134 (2006).

53. Gouin, O. et al. TRPV1 and TRPA1 in cutaneous neurogenic and chronic inflammation: pro-inflammatory response induced by their activation and their sensitization. Protein Cell (2017) doi:10.1007/s13238-017-0395-5.

54. Meng, J., Wang, J., Lawrence, G. & Dolly, J. O. Synaptobrevin I mediates exocytosis of CGRP from sensory neurons and inhibition by botulinum toxins reflects their anti-nociceptive potential. J. Cell Sci. 120, 2864–2874 (2007).

55. Huang, L.-Y. M. & Neher, E. Ca2+-Dependent Exocytosis in the Somata of Dorsal Root Ganglion Neurons. Neuron 17, 135–145 (1996).

56. Fuchs, P. N., Meyer, R. A. & Raja, S. N. Heat, but not mechanical hyperalgesia, following adrenergic injections in normal human skin. Pain 90, 15–23 (2001).

57. Ebbinghaus, M. et al. The role of interleukin-1 β in arthritic pain: main involvement in thermal, but not mechanical, hyperalgesia in rat antigen-induced arthritis. Arthritis Rheum. 64, 3897–3907 (2012).

58. Usoskin, D. et al. Unbiased classification of sensory neuron types by large-scale single-cell RNA sequencing. Nat. Neurosci. 18, 145–153 (2014).

59. Ghitani, N. et al. Specialized Mechanosensory Nociceptors Mediating Rapid Responses to Hair Pull. Neuron 95, 944–954.e4 (2017).

60. Li, C.-L. et al. Somatosensory neuron types identified by high-coverage single-cell RNA-sequencing and functional heterogeneity. Cell Res. 26, 83–102 (2016).

61. Liu, Y. & Ma, Q. Generation of somatic sensory neuron diversity and implications on sensory coding. Curr. Opin. Neurobiol. 21, 52–60 (2011).

62. Sher, E. et al. Metabolism and trafficking of N-type voltage-operated calcium channels in neurosecretory cells. J. Bioenerg. Biomembr. 30, 399–407 (1998).

63. Waterman, S. A. Role of N-, P-and Q-type voltage-gated calcium channels in transmitter release from sympathetic neurones in the mouse isolated vas deferens. Br. J. Pharmacol. 120, 393–398 (1997).

64. Burnstock, G. Purinergic Mechanisms and Pain. in Advances in Pharmacology vol. 75 91–137 (Elsevier, 2016).

65. Fields, R. D. & Burnstock, G. Purinergic signalling in neuron-glia interactions. Nat. Rev. Neurosci. 7, 423–436 (2006).

66. Pinho-Ribeiro, F. A., Verri, W. A. & Chiu, I. M. Nociceptor Sensory Neuron-Immune Interactions in Pain and Inflammation. Trends Immunol. 38, 5–19 (2017).

67. Saegusa, H. & Tanabe, T. N-type voltage-dependent Ca2+ channel in non-excitable microglial cells in mice is involved in the pathophysiology of neuropathic pain. Biochem. Biophys. Res. Commun. 450, 142–147 (2014).

68. Zeilhofer, H. U., Benke, D. & Yevenes, G. E. Chronic Pain States: Pharmacological Strategies to Restore Diminished Inhibitory Spinal Pain Control. Annu. Rev. Pharmacol. Toxicol. 52, 111–133 (2012).

69. Baddack, U. et al. Suppression of Peripheral Pain by Blockade of Voltage-Gated Calcium 2.2 Channels in Nociceptors Induces RANKL and Impairs Recovery From Inflammatory Arthritis in a Mouse Model: AGGRAVATION OF ARTHRITIS BY ANALGESIC CONOTOXINS. Arthritis Rheumatol. 67, 1657–1667 (2015).

70. Miljanich, G. P. Ziconotide: neuronal calcium channel blocker for treating severe chronic pain. Curr. Med. Chem. 11, 3029–3040 (2004).

71. Staal, J. B., de Bie, R. A., de Vet, H. C. W., Hildebrandt, J. & Nelemans, P. Injection Therapy for Subacute and Chronic Low Back Pain: An Updated Cochrane Review. Spine 34, 49–59 (2009).

72. Williams, J. A., Day, M. & Heavner, J. E. Ziconotide: an update and review. Expert Opin. Pharmacother. 9, 1575–1583 (2008).

73. Marangoudakis, S. et al. Differential Ubiquitination and Proteasome Regulation of Ca_V_2.2 N-Type Channel Splice Isoforms. J. Neurosci. 32, 10365–10369 (2012).

74. Cavanaugh, D. J. et al. Trpv1 Reporter Mice Reveal Highly Restricted Brain Distribution and Functional Expression in Arteriolar Smooth Muscle Cells. J. Neurosci. 31, 5067–5077 (2011).

75. Madisen, L. et al. A toolbox of Cre-dependent optogenetic transgenic mice for light-induced activation and silencing. Nat. Neurosci. 15, 793–802 (2012).

76. Madisen, L. et al. A robust and high-throughput Cre reporting and characterization system for the whole mouse brain. Nat. Neurosci. 13, 133–140 (2010).

77. Maccarrone, M. Characterization of the endocannabinoid system in boar spermatozoa and implications for sperm capacitation and acrosome reaction. J. Cell Sci. 118, 4393–4404 (2005).

78. Mizrak, S. C. & van Dissel-Emiliani, F. M. F. Transient receptor potential vanilloid receptor-1 confers heat resistance to male germ cells. Fertil. Steril. 90, 1290–1293 (2008).

79. Francavilla, F. et al. Characterization of the endocannabinoid system in human spermatozoa and involvement of transient receptor potential vanilloid 1 receptor in their fertilizing ability. Endocrinology 150, 4692–4700 (2009).

80. Hargreaves, K., Dubner, R., Brown, F., Flores, C. & Joris, J. A new and sensitive method for measuring thermal nociception in cutaneous hyperalgesia: Pain 32, 77–88 (1988).

